# A role for Hypoxia Inducible Factor 1a (HIF1a) in intermittent hypoxia-dependent changes to spatial memory and synaptic plasticity

**DOI:** 10.1101/595975

**Authors:** Alejandra Arias-Cavieres, Maggie A. Khuu, Chinwendu U Nwakudu, Jasmine E. Barnard, Gokhan Dalgin, Alfredo J. Garcia

**Author notes:** **Footnote:** This article was first published as a preprint: A Arias-Cavieres, MA Khuu, CU Nwakudu, JE Barnard, G Dalgin, AJ Garcia III (2019). *A role for Hypoxia Inducible Factor 1a (HIF1a) to intermittent hypoxia-dependent changes in spatial memory and synaptic plasticity.* bioRxiv. https://doi.org/10.1101/595975.

## Abstract

Intermittent hypoxia (IH), a key feature of sleep apnea, increases the oxygen regulated transcription factor Hypoxia Inducible Factor 1a (HIF1a). Although recognized for its role in IH-dependent changes in cardio-respiratory physiology, it remains unclear how IH-dependent HIF1a signaling affects neurophysiology underlying learning and memory. This study examines how IH affects hippocampal associated learning and memory in wildtype mice and mice heterozygous for the HIF1a gene (HIF1a^+/-^). In wild-type mice, ten days of IH impaired performance in the Barnes maze increased hippocampal HIF1a and elevated protein carbonyls. These behavioral and biochemical effects of IH were accompanied by a decrease in the N-Methyl-D-Aspartate receptor (NMDAr) and an attenuation of long-term potentiation (LTP) in area CA1. In HIF1a^+/-^, IH did not impair Barnes maze performance, increase hippocampal HIF1a, or enhance protein carbonyl content. At the network level, IH neither led to a decrease in NMDAr nor impaired LTP. Concurrent antioxidant treatment during IH mitigated the IH-dependent effects on the Barnes maze performance and LTP in wildtype mice. Our findings indicate that IH-dependent HIF1a signaling leads to oxidative stress and reduces NMDAr to impair LTP in area CA1, which contributes to IH-dependent deficits in learning and memory associated with the hippocampus.

**Significance:** Intermittent Hypoxia is a hallmark of sleep apnea and decreases the threshold for cognitive deficit. We demonstrate that intermittent hypoxia-dependent HIF1a signaling contributes to impairments in hippocampal associated memory. This is co-incidental with HIF1a-mediated alternations in synaptic physiology and increased oxidative stress.

**Key points:**

- Intermittent hypoxia (IH) is a hallmark of sleep apnea and is known to cause learning and memory deficits.
- Hypoxia Inducible Factor 1a (HIF1a), is associated with IH-dependent changes in physiology.
- IH exposure causes increased hippocampal HIF1a in wild type mice and is associated with elevated oxidative stress, impairments to spatial memory, and suppression of long term potentiation (LTP).
- IH-dependent suppression of LTP is co-incidental with diminished NMDA receptor contribution to glutamatergic transmission.
- Following IH, mice heterozygous for HIF1a (HIF1a^+/-^) do not show an increase in HIF1a and oxidative stress, or changes in either behavior or glutamatergic transmission.

## Introduction

While sleep apnea is recognized as a risk factor for cardiorespiratory dysfunction, this condition also decreases the threshold for cognitive impairment (Devita et al., 2017; Leng et al., 2017). Similarly, intermittent hypoxia (IH), a hallmark of sleep apnea, has been shown to cause both autonomic dysfunction (Fang and Chen, 1973; Lesske et al., 1997; Greenberg et al., 1999) and memory deficits (Wallace and Bucks, 2013). At the cellular level, exposure to IH increases hypoxia inducible factor 1a (HIF1a), a ubiquitous transcription factor regulated by the state of oxygenation. Although commonly recognized as a crucial mediator of cellular adaptations to hypoxia, HIF1a has been implicated as a critical factor facilitating maladaptive changes in cardiorespiratory physiology caused by IH (Prabhakar, 2016). However, the role of HIF1a signaling in IH-dependent changes to the neurophysiology remains poorly understood.

The hippocampal formation is important for learning and memory (Morris et al., 1982) and has been frequently identified as a brain region affected in sleep apnea (Kuwabara et al., 2009; Encinas et al., 2011; Licht et al., 2011; Macey et al., 2018; Huang et al., 2019; Owen et al., 2019). Biochemical studies have documented IH-dependent increases in hippocampal HIF1a protein (Chou et al., 2013; Wall et al., 2014) and oxidative stress within the brain (Nair et al., 2011; Chou et al., 2013). Neurophysiological investigations have also documented that IH exposure leads to deficits in spatial learning and memory (Row et al., 2002; Row et al., 2003) which coincide with weakened hippocampal LTP in area CA1 (Gozal et al., 2002; Goldbart et al., 2003; Payne et al., 2004; Xie et al., 2010; Zhang et al., 2012; Wall et al., 2014). HIF1a signaling may have an important protective pro-survival role in the brain preserving function in response to the hypoxia experienced during IH. Alternatively, HIF1a may serve as a pro-oxidant transcription factor leading to oxidative stress and impaired neurophysiology.

To better resolve the role of IH-dependent increases in HIF1a, we examine how IH affects behavior and synaptic physiology in wildtype mice and mice heterozygous for HIF1a (HIF1a^+/-^). Impaired performance on the Barnes maze was evident following IH. This phenotype was co-incidental with increased hippocampal oxidative stress, a decreased contribution of the NMDA receptor (NMDAr) to the field excitatory postsynaptic potential (fEPSP), and suppression of LTP. In contrast, IH neither impaired behavioral performance nor caused hippocampal oxidative stress in HIF1a^+/-^. HIF1a^+/-^ exposed to IH did exhibit a decrease in NMDAr contribution to the field excitatory postsynaptic potential (fEPSP) or show an attenuation in LTP. These findings indicate that enhanced HIF1a signaling is a significant factor contributing to IH-dependent changes in synaptic physiology and spatial learning and memory associated with the hippocampus.

## Methods

### Study Approval

In accordance with National Institutes of Health guidelines, all animal protocols were performed with approval from the Institute of Animal Care and Use Committee at The University of Chicago.

### Animals

Animals were housed in AAALAC-approved facilities with a 12 hour/12 hour light-dark cycle and given ad libitum access to food and water. Experiments were performed on wildtype mice and HIF1a^+/-^ (Iyer et al., 1998), (Peng et al., 2006) from both sexes (P50 to P70). All animals were maintained on a C57BL/6 background. Automated genotyping was performed independently by a commercial service (Transnetyx Inc).

### Intermittent hypoxia (IH) exposure

Male and female mice (P60-P80) were exposed to chronic intermittent hypoxia for ten consecutive days (IH_10_). In brief, as previously described (Peng et al., 2003), the IH_10_ paradigm was performed in a special chamber during the light cycle and lasted 8 hours per day (i.e., 80 intermittent hypoxia cycles/day). A single hypoxic cycle was achieved by flowing 100% N_2_ into the chamber for approximately 60s (nadir O2 reached 4.5±1.5% and followed immediately by an air break (∼21% O_2_; 300s).

In a subset of animals, manganese (III) tetrakis(1-methyl-4-pyridyl) porphyrin (MnTMPyP, Enzo Life Sciences, CAT #: ALX-430-070), was administered via intraperitoneal injection at the beginning of each day prior to exposure to IH. Previous reports have indicated that dose of MnTMPyP at either 5mg/kg (Peng et al., 2013) or 15mg/kg (Khuu et al., 2019) can mitigate the effects of IH in the nervous system. Therefore, the smaller dose (5 mg/kg, n=3 mice) and the larger dose (15mg/kg, n=4 mice) were used but no differences were evident between dosage groups; and therefore, the data at the two concentrations were pooled.

### Barnes maze

The Barnes maze was performed using a custom made opaque white circular acrylic platform (92.4 cm in diameter) with 20 equidistant holes (5.08 cm in diameter and 2.54 cm from the edge). The platform was elevated (30 cm from the floor) ground and surrounded by four identical walls (27.94 cm high). By default, each hole was closed with a fixed piece of opaque acrylic that could be removed to lead to a dark exit box. Lighting was achieved through diffuse overhead fluorescent lighting such that all holes were equally lit. An overhead camera was suspended above the maze. Data collection and *posthoc* analysis was performed using CinePlex Video Tracking System (Plexon, Dallas, TX).

As previously described (Christakis et al., 2012), the task was performed using a four-day protocol consisting of one training trial per day for three consecutive days and a probe trial on the fourth day. Barnes Maze began on the seventh day of IH_10_ exposure with respective controls run at the same time. For the training trials, all but one of the holes (exit hole) was closed. An exit box with a small ramp was placed directly underneath the exit hole. Animals were given a maximum of 6 minutes to locate the exit and if unable to locate the exit, they were gently guided to the exit. If the mouse found and entered the exit before the six minutes was over, the trial ended at the time that the mouse entered the exit and the mouse was promptly returned to its home cage. During the probe trial, all holes were closed, and the animal was given 6 minutes to explore the maze. The entire arena was sanitized in-between trials. Following the end of behavior, IH animals were immediately returned to the IH chamber. For animals in the MnTMPyP group, MnTMPyP was administered after the daily behavioral testing, but prior to the return to the IH chamber.

### Slice Preparation

As previously described (Khuu et al., 2019), acute coronal hippocampal slices were prepared from mice unexposed to intermittent hypoxia (control) or from mice exposed to IH for ten days (IH_10_). Tissue harvest occurred within one to two days following IH_10_. Mice were anesthetized with isoflurane and euthanized via rapid decapitation. The cerebrum was immediately harvested and blocked, rinsed with cold artificial cerebrospinal fluid (aCSF), and mounted for vibratome sectioning. The mounted brain tissue was submerged in aCSF (4°C; equilibrated with 95% O_2_, 5% CO_2_) and coronal cortico-hippocampal brain slices (350 µm thick) were prepared. Slices were then immediately transferred into a holding chamber containing aCSF equilibrated with 95% O_2_, 5% CO_2_ (at 20.5±1°C). Slices were allowed to recover for a minimum of one hour prior to recording and used up to eight hours following tissue harvest. The composition of aCSF (in mM): 118 NaCl, 10 glucose, 20 sucrose, 25 NaHCO_3_, 3.0 KCl, 1.5 CaCl_2_, 1.0 NaH_2_PO_4_ and 1.0 MgCl_2._

### Extracellular recording of the field excitatory postsynaptic potential

For electrophysiological recordings, slices were transferred to a recording chamber with recirculating aCSF (30.5±1°C, equilibrated with 95% O_2_ and 5% CO_2_) and allowed 15 min to acclimate to the recording environment. The fEPSP in the CA1 was evoked by electrical stimulation. The stimulation electrode was positioned in Schaffer Collateral and recording electrode (1-2 MΩ) was placed into the stratum radiatum of the CA1. The intensity of the electrical current (100-400 µA; 0.1-0.2 ms duration) was set to the minimum amount of current required to generate ∼50 % of the maximal initial slope (m_i_) of the fEPSP. The current stimulus to evoked fEPSP relationship was examined across range of stimuli between 0 to 700 microamperes (µA), sampled at intervals of 50 µA 0.2ms duration. This relationship was examined in aCSF, Mg^2+^ free aCSF, and Mg^2+^ free aCSF with 20 µM APV (Sigma-Aldrich, MN). The composition of Mg2+ free aCSF (in mM): 119.5 NaCl, 10 glucose, 20 sucrose, 25 NaHCO_3_, 3.0 KCl, 1.5 CaCl_2_, 1.0 NaH_2_PO_4_. The NaCl was increased to 119.5 mM to keep osmolarity from changing when switching from aCSF to Mg^2+^ free aCSF. The fEPSP was evoked every 20 s. After 10 minutes of recording the baseline fEPSP, LTP was induced using high frequency stimulation (HFS). HFS consisted four 500 msec trains of stimuli (100 Hz) given at 30 sec intervals. The fEPSP slope was normalized to baseline values (prior to HFS).

All recordings were made using the Multicamp 700B (Molecular Devices: https://www.moleculardevices.com/systems/conventional-patch-clamp/multiclamp-700b-microelectrode-amplifier). Acquisition and post hoc analyses were performed using the Axon pCLAMP10 software suite (Molecular Devices: https://www.moleculardevices.com/system/axon-conventional-patch-clamp/pclamp-11-software-suite).

### Western Blot

Western blot assays were performed using hippocampal tissue homogenates for quantitative analysis of HIF1a (R and D Systems Cat# AF1935, RRID:AB_355064) and PCNA (Bethyl Cat# A300-276A, RRID:AB_263393) content. Stepwise separation of cytoplasmic and nuclear protein extracts were prepared by NE-PER nuclear and cytoplasmic extraction kit (Thermo Scientific, 78833) by following manufactures instructions. Briefly, cytoplasmic fragment was obtained by homogenizing tissue using a tissue grinder and then by pipetting in cytoplasmic extraction buffers. After isolation of cytoplasmic fragment, the insoluble pellet that contains nuclear proteins was suspended in nuclear extraction buffer and separated by centrifugation. Halt Protease Inhibitor (Thermo Scientific, 1860932) was added into cytoplasmic and nuclear extraction buffers to prevent protein degradation. All analyses were conducted by Raybiotech, Inc. (Norcross, GA), using the automated Capillary Electrophoresis Immunoassay machine (WES™, ProteinSimple Santa Clara, CA). The samples, blocking reagent, wash buffer, primary antibodies, secondary antibodies, and chemiluminescent substrate were dispensed into designated wells in the manufacturer provided microplate. After plate loading, the separation electrophoresis and immunodetection steps took place in the capillary system and were fully automated. Auto Western analysis was carried out at room temperature, and instrument default settings were used.

### Protein Carbonyls

Whole cell protein lysates were isolated from hippocampal tissues by using M-PER mammalian protein extraction reagent (Thermo Scientific, 78501) and by adding Halt Protease Inhibitor (Thermo Scientific, 1860932). Protein lysates were immediately processed or kept in −80°C until used. The amount of protein carbonyls was determined using a Protein Carbonyl Colorimetric Assay Kit (Cayman Chemical, Cat#10005020), per manufacturer instructions and absorbance was measured at a wavelength between 360-385 nm using a plate reader. Protein content was determined using a Protein Determination Kit (Cayman Chemical, Cat# 704002) and absorbance was measured at 595 nm using a Nanodrop 2000 spectrophotometer (Thermofisher, Cat# ND-2000).

### Experimental Design and Statistical Analyses

All n values are total number of animals, unless otherwise noted. Statistics were performed using Origin 8 Pro (OriginLab, RRID:SCR_014212) or Prism 6 (GraphPad Software; RRID:SCR_015807). Comparisons between two groups were conducted using unpaired two-tailed t tests with Welch’s correction. To compare three groups, a one-way ANOVA was performed followed by a posthoc Tukey’s multiple comparison test or Dunnett’s test, as appropriate. To assess the fEPSP across a range of stimuli, a two-way ANOVA was performed followed by a Bonferroni post-test for significance. Results are presented as mean ± S.E.M. were considered significant when the P-value was less than 0.05.

## Results

HIF1a protein content was measured in nuclear extracts prepared from wildtype hippocampi unexposed to IH (control) and wildtype hippocampi exposed to ten days of IH (IH_10_). Nuclear HIF1a was approximately two times greater in extracts from IH_10_ when compared to control (**Figure 1A**, control n=5, IH_10_ n=4, P=0.019). In contrast, nuclear hippocampal HIF1a was unchanged in extracts from hippocampi of HIF1a^+/-^ mice unexposed to IH (0-HIF1a^+/-^) when compared to tissue originating from HIF1a^+/-^ mice exposed to ten days of IH (10-HIF1a^+/-^) (**Figure 1B**, 0-HIF1a^+/-^, n=4, 10-HIF1a^+/-^ n=4, P=0.84). These findings demonstrate that IH-dependent increases in HIF1a protein are mitigated in mice possessing a single copy of the gene.

**Figure 1.**
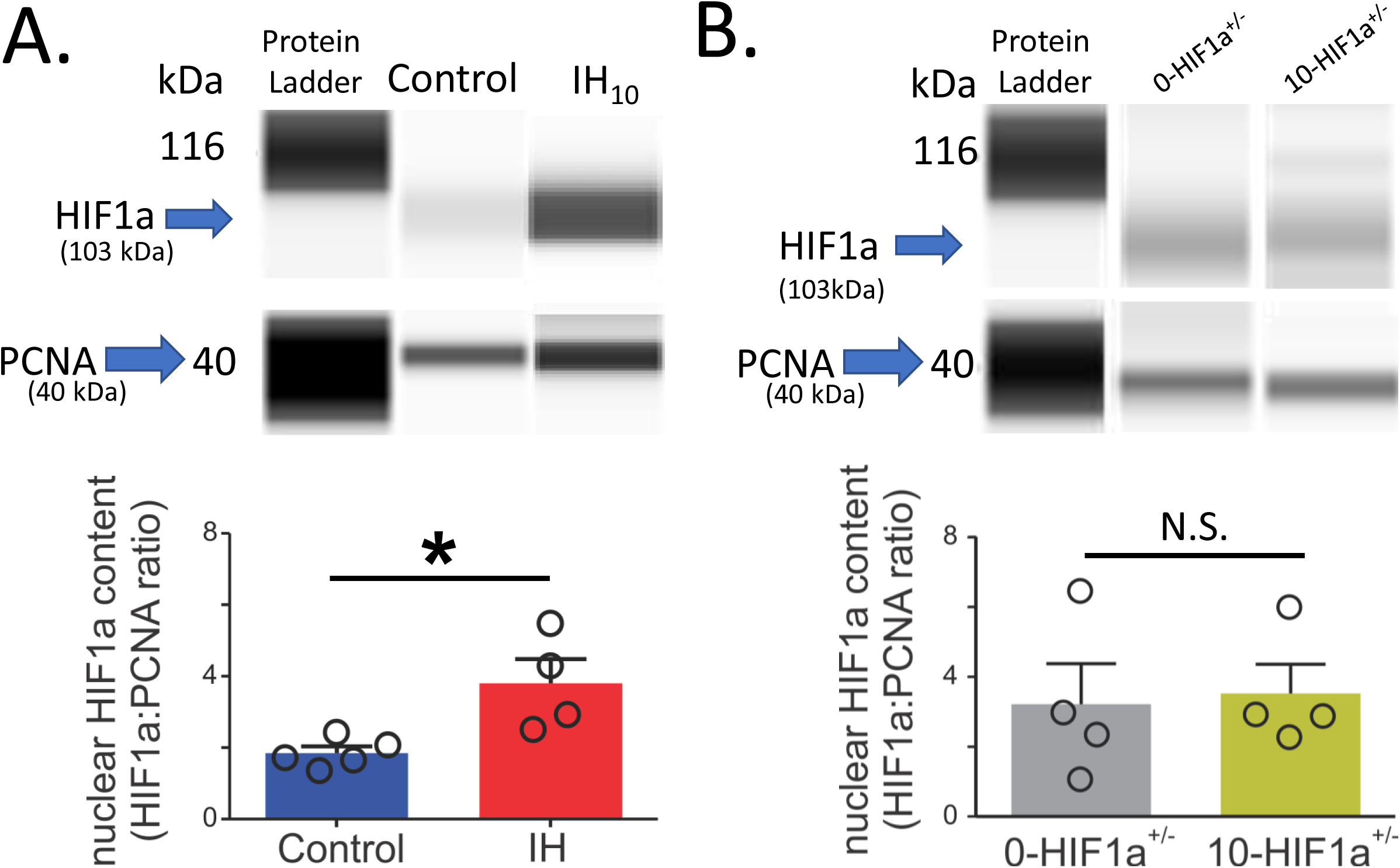
Ten days of IH increases hippocampal hypoxia inducible factor 1a (HIF1a) in wild type mice but not in HIF1a+/- mice. **A.** *top:* Representative digitized western blot images for HIF1a (103 kDa) and Proliferating Cell Nuclear Antigen (PCNA, 40 kDa) in hippocampal nuclear protein fractions from control (n=5) and IH_10_ (n=4). *bottom:* Quantification of HIF1a protein normalized to PCNA revealed that nuclear HIF1a was increased in IH_10_ when compared to control (P=0.019). **B.** *top:* Representative digitized western blot images HIF1a and PCNA in hippocampal nuclear protein fractions from 0-HIF1a^+/-^ (n=4) and 10-HIF1a^+/-^ (n=4). *bottom:* Quantification of HIF1a protein normalized to PCNA revealed that nuclear HIF1a was not different between 0-HIF1a^+/-^ and 10-HIF1a^+/-^ (P=0.84).

To determine the behavioral consequences of IH on wildtype mice and HIF1a^+/-^, we examined spatial learning and memory by assessing performance in a Barnes maze apparatus for four experimental groups: wildtype unexposed mice (control, n=11), wildtype mice exposed to ten days of IH (IH_10_, n=10), HIF1a+/- mice unexposed to IH (0-HIF1a^+/-^, n=7), and HIF1a^+/-^ mice exposed to ten days of IH (10-HIF1a^+/-^, n=8). During training, control and IH_10_ exhibited progressive improvement on locating the exit zone as indicated by the decrease in latency to exit over course of three training sessions (**Figure 2A and 2B**). 0-HIF1a^+/-^ and 10-HIF1a^+/-^ groups displayed similar performance during training (**Figure 2C and 2D**). During the probe trial (when the exit was closed), performance of each group was assessed by examining: 1) distance travelled to the exit zone; 2) primary latency to exit zone entry; and 3) probability to exit zone entry.

**Figure 2.**
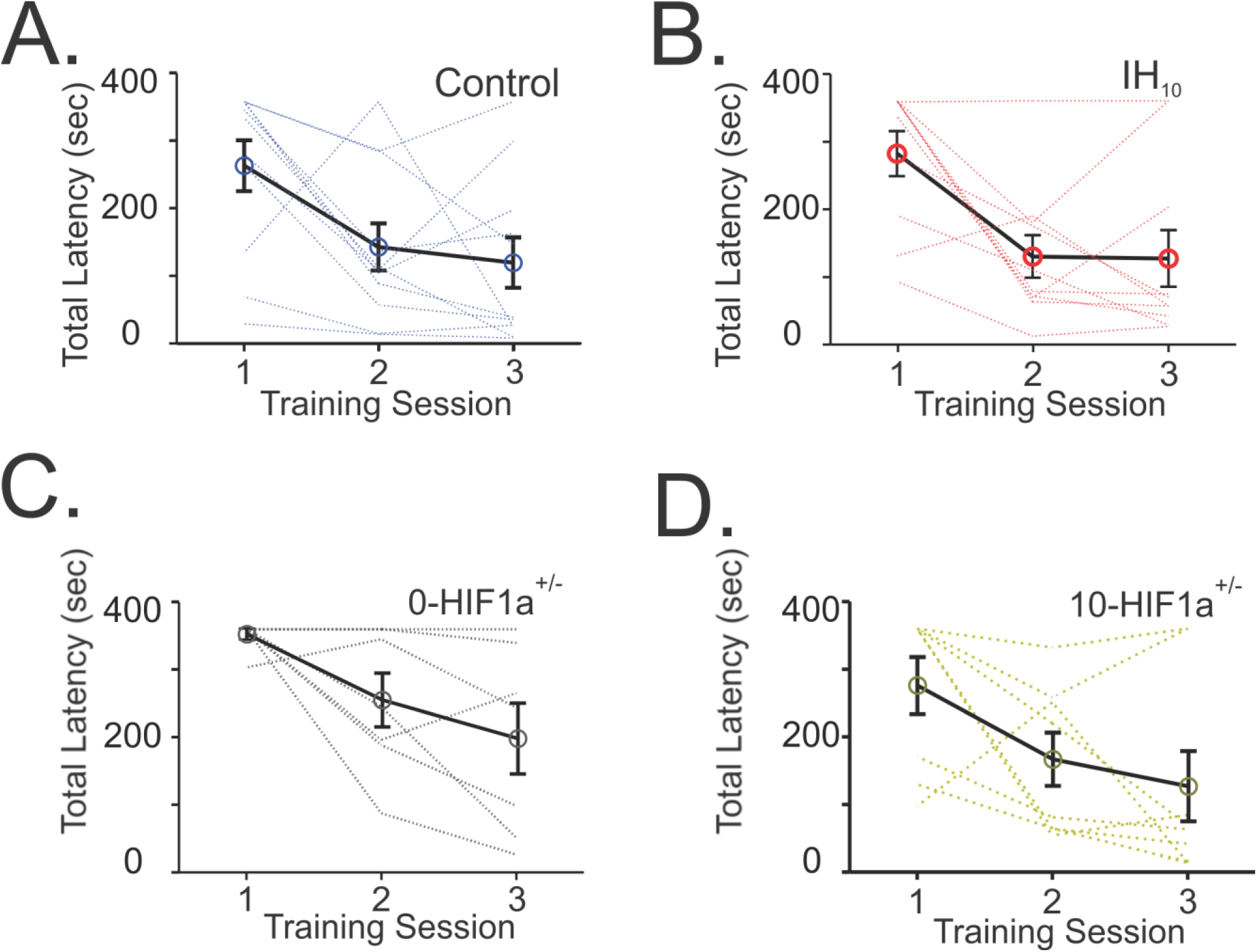
Barnes maze training performance in control, IH_10_, 0-HIF1a+/-, and 10-HIF1a+/-. **A.** Control (n=10), each blue line represents an individual performance during training; **B.** IH_10_ (n=11), each red line represents an individual performance during training; **C.** 0-HIF1a^+/-^ (n=7), each gray line represents an individual performance during training; and **D.** 10-HIF1a^+/-^ (n=8), each yellow line represents an individual performance during training. Training to the exit was conducted over three sessions. Each session was separated by 24 hours. All experimental groups exhibit decreased total latency over the course of training.

The distance travelled to the exit zone was greater in IH_10_ when compared to control (control: 2.60 ± 0.70 m versus IH_10_: 10.34 ± 3.32m, P=0.048). Similarly, larger primary latency to exit zone entry was observed in IH_10_ (**Figure 3B**, control: 22.60 ± 6.28 sec versus IH_10_: 117.90 ± 37.47m, P=0.034). In contrast, 0-HIF1a+/- mice to 10-HIF1a^+/^ mice were similar in distance travelled to the exit zone (**Figure 3C**, 0-HIF1a^+/-^=2.37 ± 0.91 m, 10-HIF1a^+/-^=1.71 ± 0.50 m; P=0.55), and primary latency to exit zone entry (**Figure 3D**, 0-HIF1a^+/-^=35.18 ± 12.28 sec, 10-HIF1a^+/-^=: 61.61 ± 26.30 sec; P=0.39).

**Figure 3.**
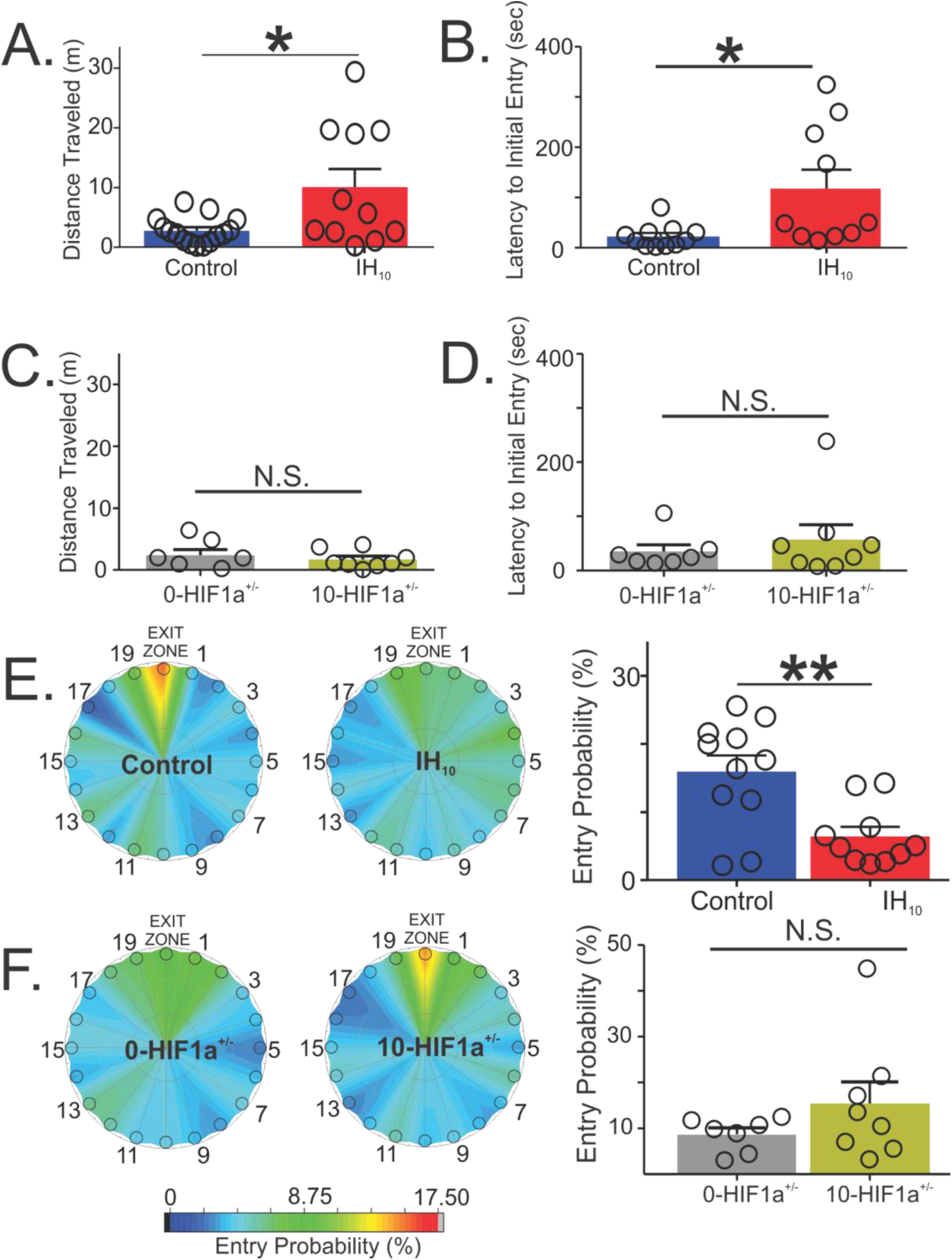
Differences in Barnes maze performance during the probe trial are apparent in wildtype mice exposed to IH but not in 10-HIF1a+/-. **A.** During the probe trial, the distance traveled to initially enter the exit zone was shorter in control (n=10) when compared to IH_10_ (n=11, P=0.048). **B**. Latency to initial entry was also smaller in control as well (P=0.034). **C**. In HIF1a^+/-^, the primary distance traveled was similar between 0-HIF1a^+/-^ and 10-HIF1a^+/-^ (P=0.55). **D**. Primary Latency between 0-HIF1a^+/-^ and 10-HIF1a+/- were similar (P=0.39). **E**. Heat maps of the entry probability across all false exits (1 to 19) and the exit zone during the probe trial illustrates a higher probability for entering the exit zone in control (left) as compared IH_10_ (center). Comparison of entry probability into the exit zone during the probe trial (right) reveals that control has a greater probability for entering the exit zone when compared to IH_10_ (P=0.004). **F**. Heat maps of the entry probability into the exit zone during the probe trial illustrates that 0-HIF1a^+/-^ (left) and 10-HIF1a^+/-^ (center) have similar probabilities for entering the exit zone. This is evident in the comparison between groups (right) (P=0.21).

Examining the probability to exit zone entry demonstrated that control consistently discriminated the location of exit hole against the other holes (**Figure 3E left**). However, in IH_10_, this was not apparent (**Figure 3E center**). These observations reflected in a difference between control and IH_10_ (**Figure 3E right**; control: 15.93 ± 2.39% versus IH_10_: 6.44 ± 1.38%, P=0.004). In contrast, entry probability into the exit zone for 0-HIF1a+/- and 10-HIF1a^+/^ was similar between both groups (**Figure 3F**, 0-HIF1a^+/-^=8.75 ± 1.38%, 10-HIF1a^+/-^=15.51 ± 4.73% m; P=0.21).

The IH-dependent differences in Barnes maze performance may be related to potential differences in synaptic physiology. To address this possibility, we characterized unpotentiated and potentiated synaptic transmission in hippocampal brain slices from the four experimental groups. In slices from wildtype mice, IH did not impact the maximal fiber volley (control= 0.32 ± 0.12 mV, n=4; IH_10_= 0.34 ± 0.08 mV, n=5; P=0.90) or the maximal unpotentiated fEPSP (control= 1.21 ± 0.20 mV, n=8; IH_10_= 1.03 ± 0.11 mV, n=10; P=0.41). No differences between groups were found in the stimulus intensity current to fiber volley relationship in aCSF, in Mg^2+^-free media, or in Mg^2+^-free media with the competitive NMDAr blocker, APV (*data not shown*). Comparing the evoked fEPSP at 700 μA revealed that synaptic transmission from control and IH_10_ were similar in aCSF (**Figure 4A**, control n=5, IH_10_ n=8, comparison at 700μA: P>0.05) and in Mg2+-free media (**Figure 4B**, comparison at 700μA: P>0.05). However, in APV, fEPSP amplitude was larger in IH_10_ (**Figure 4C**, comparison at 700μA: P<0.05), indicating contribution of ionotropic glutamate receptors to the fEPSP change following IH. In slices from HIF1a^+/-^ animals, the maximal fiber volley (0-HIF1a^+/-^= 0.32 ± 0.12 mV, n=4; 10-HIF1a^+/-^= 0.34 ± 0.08 mV, n=5; P=0.90) as well as the maximal fEPSP (0-HIF1a^+/-^= 0.63 ± 0.06 mV, n=7; 10-HIF1a^+/-^= 1.13 ± 0.11 mV, n=8; P=0.0003) were consistently smaller in 0-HIF1a^+/-^ when compared to 10-HIF1a^+/-^. The fiber volley was undetectable when evoked sub-maximally; therefore, we were unable to compare the stimulus intensity-fiber volley relationship between 0-HIF1a^+/-^ and 10-HIF1a^+/-^. Differences in the stimulus intensity-fEPSP relationship were present between 0-HIF1a^+/-^ and 10-HIF1a^+/-^ in aCSF (**Figure 4D**, comparison at 700μA: P<0.05) as well as Mg^2+^-free media (**Figure 4E**, comparison at 700μA: P<0.05) but were no longer present in APV (**Figure 4F**, comparison at 700μA: P>0.05).

**Figure 4.**
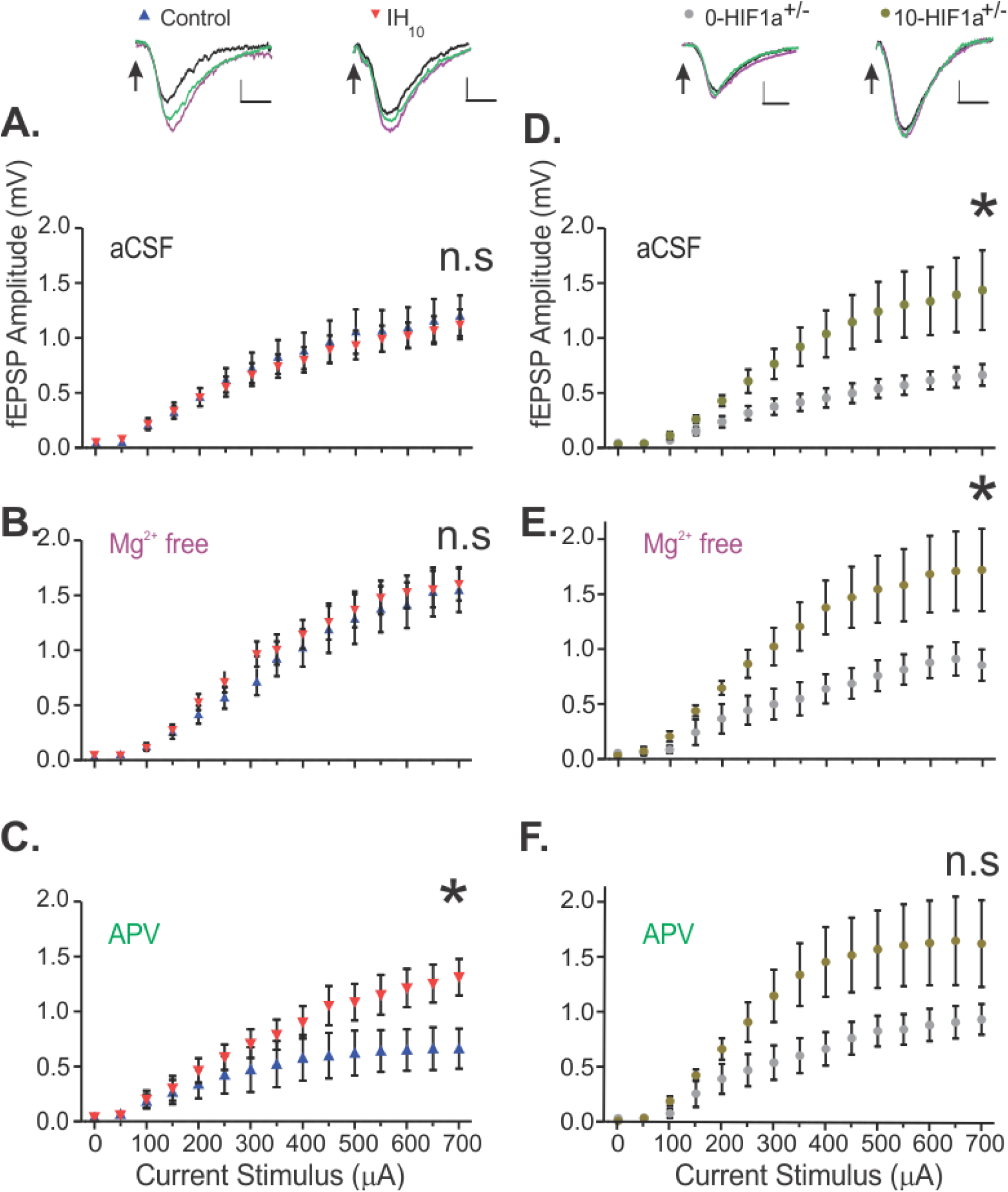
Examination of the current stimulus-fEPSP relationship reveals that IH reduces NMDAr mediated reductions in synaptic transmission in area CA1 of wildtype but not in HIF1a+/-. **A.** No differences were found in the current stimulus-fEPSP amplitude relationship between control (n=5) and IH_10_ (n=8) groups i**n** ACSF (P>0.05). **B**. fEPSP amplitudes between control (n=5) and IH_10_ treated animals in Mg2+-free media (P>0.05) were also similar. **C**. In Mg^2+^-free media, blockade of the NMDAr with APV (20µM), reduced the fEPSP amplitude of the control to a greater degree than IH10 (P<0.05). **D**. The fEPSP of 0-HIF1a+/- (n=4) was smaller compared to 10-HIF1a^+/-^(n=7) in aCSF (P<0.05). **E**. Slices in Mg^2+^-free media also revealed that fEPSP of 0-HIF1a+/- (n=4) was smaller compared to 10-HIF1a^+/-^(n=7) (P<0.05). **F**. However, fEPSP of 0-HIF1a+/- (n=4) was not different compared to 10-HIF1a^+/-^(n=7) with Mg^2+^-free media with APV (P>0.05). Representative traces above the graphs show fEPSPs for the four representative animal groups under each of the three different medias (ACSF= black, Mg^2+^ free media = magenta, APV=green). Scale bars: 0.2 mV/10 ms. Two-way ANOVA were performed in each condition followed by a *posthoc* Bonferroni comparison. P-values reported here represent the comparison made at 700µA for each condtion. *=P<0.05; N.S.=P>0.05.

The NMDAr contributes to the prolonging of decay time of the fEPSP. Therefore, we also examined decay time of the fEPSP in Mg^2+^-free media with and without APV. The NMDAr blocker reduced the decay time by −70 ± 11% in the control group (n=5); whereas, in the IH_10_ group (n=8) APV reduced the decay time of fEPSP by −28 ± 16%. When comparing these changes, the effectiveness of APV was greater in control as compared to IH10 (**Figure 5A**, P=0.016). However, reduction in decay time of the fEPSP by APV was similar between 0-HIF1a^+/-^ (−24 ± 12%, n=7) and 10-HIF1a^+/-^ (−36 ± 11%n=8) (**Figure 5B**, P=0.46) further supporting an IH-dependent reduction in the NMDAr. These findings suggest that IH acts to reduce NMDAr through a HIF1a mediated mechanism.

**Figure 5.**
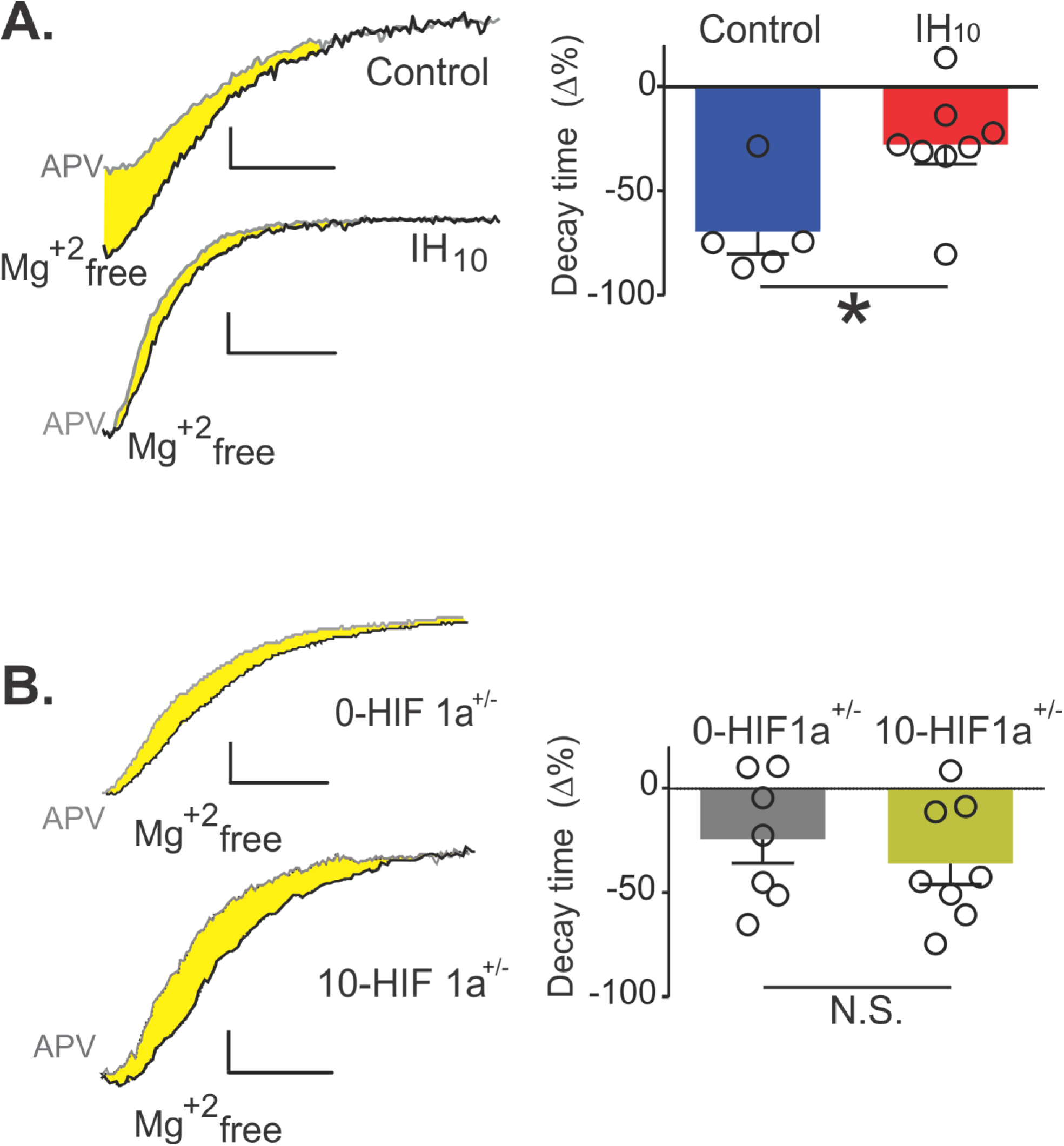
The effect of IH on decay time of the fEPSP from wildtype and HIF1a+/-. **A.** *Left:* Representative traces of the fEPSP in control and IH_10_ in Mg2+-free media and in APV (20μM). Yellow shaded region highlights differences in decay times. *Right:* The efficacy for APV to reduce decay time of the fEPSP in Mg^2+^ free media is greater in control versus IH_10_ (P=0.015). **B.** *Left:* Representative traces of fEPSP decay after in 0-HIF1a^+/-^ and 10-HIF1a^+/-^ in Mg^2+^-free media and in APV (20μM). Yellow shaded region highlights differences in decay between conditions. Scale bars: 0.2 mV/10 ms *Right:* APV has similar effects on reducing decay time of the fEPSP in Mg^2+-^ free media in 0-HIF1a^+/-^ and 10-HIF1a^+/-^ (P=0.46).

To examine the consequence that reduced NMDAr contribution may have on synaptic plasticity, we evoked LTP using HFS. HFS evoked LTP in area CA1 from both control and IH_10_ (**Figure 6A**, control n=7, IH_10_ n=11). No differences between experimental groups were observed in the fEPSP immediately following post-tetanic stimulation (**Figure 6B**, control=129±9**%** versus IH_10_=117±3% P=0.26). However, at 60 min following HFS, the magnitude of LTP in the control versus the IH_10_ group was larger (**Figure 6C**, control=162±12% IH_10_=122±7%, P=0.015). In the HIF1a^+/-^ group, HFS evoked LTP in 0-HIF1a^+/-^ (n=4) and in 10-HIF1a^+/-^ (n=6) was similar (**Figure 6D**). The strength of potentiation in 0-HIF1a^+/-^ and in 10-HIF1a^+/-^ was comparable immediately following HFS (**Figure 6E**, 0-HIF1a^+/-^=117 ± 6.72%, 10-HIF1a^+/-^=125 ±12.15%, P=0.59), and 60 min following HFS (**Figure 6F**, 0-HIF1a^+/-^=137 ± 21.47%, 10-HIF1a^+/-^=151.9 ± 9.58%, P=0.56).

**Figure 6.**
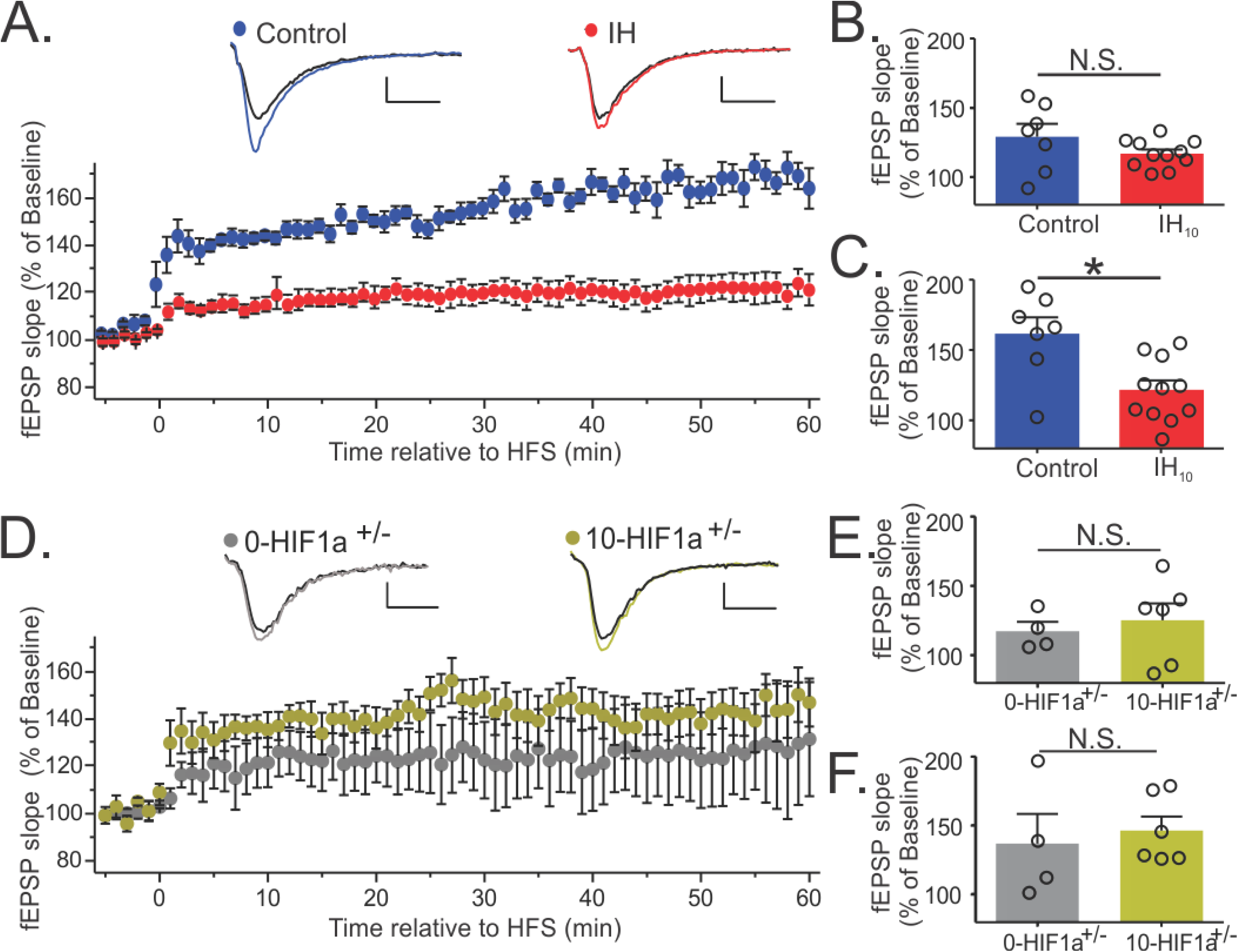
IH suppresses LTP in area CA1 of the hippocampus. **A.** LTP was evoked using high frequency stimulation (HFS) in control (blue, n=7) and IH_10_ (red, n=11). **B.** No difference between control and IH_10_ was observed in the slope of the fEPSP immediately following HFS (P=0.26). **C.** 60 min following HFS the magnitude of potentiation in control was larger than IH_10_ (P=0.015). **D**. LTP was evoked in both groups 0-HIF1a^+/-^ and 10-HIF1a^+/-^. **E**. No significant difference was found between 0-HIF1a^+/-^ and 10-HIF1a^+/-^ (P=0.59) at 10 min HFS. F. At 60 min after HFS both groups don’t show significant difference (P=0.56). Representative traces illustrate baseline (black) and 60 mins following HFS (colored trace). Scale bars 0.2 mV/10 ms.

We next sought to determine whether IH caused increased oxidative stress by measuring protein carbonyl content in hippocampal homogenates from control (n=4), IH_10_ (n=4), 0-HIF1a+/- (n=4), and 10-HIF1a+/- (n=4). Relative to the control, protein carbonyl content was elevated in IH_10_ (**Figure 7**, P<0.01) but protein carbonyl content was not changed in homogenates from either 0-HIF1a^+/-^ or 10-HIF1a^+/-^ (**Figure 7**). These findings indicate that hippocampal oxidative stress is a downstream consequence of enhanced HIF1a signaling due to IH exposure. To determine the involvement of oxidative stress in the Barnes Maze performance and synaptic of physiology in area CA1, wildtype mice treated with the antioxidant MnTMPyP, for 10 days without IH exposure (0-MnTMPyP, n=9) and during exposure to 10 days of IH (10-MnTMPyP n=7). Both groups exhibited improvement in exiting the maze with training (**Figure 8A**). During the probe trial, neither latency to first entry (0-MnMTPyP = 24 ± 7sec, 10MnTMPyP = 38±15sec; P=0.42) nor path length (0-MnMTPyP = 0.25 ± 0.08m, 10-MnTMPyP = 0.29 ± 0.05 sec; P=0.70) was different between 0-MnTMPyP and 10-MnTMP. Additionally, the probability of entering the exit zone during the probe trial was similar between groups (**Figure 8A**). In hippocampal slices from 10-MnTMPyP (n=8), LTP was readily evoked (**Figure 8B**). However, when comparing the magnitude of LTP in control, IH_10_ and 10-MnTMPyP revealed that the magnitude of LTP in 10-MnTMPyP was neither different when compared to control or to IH_10_ (**Figure 8C**, One-ANOVA followed by a Tukey’s multiple comparisons test).

**Figure 7.**
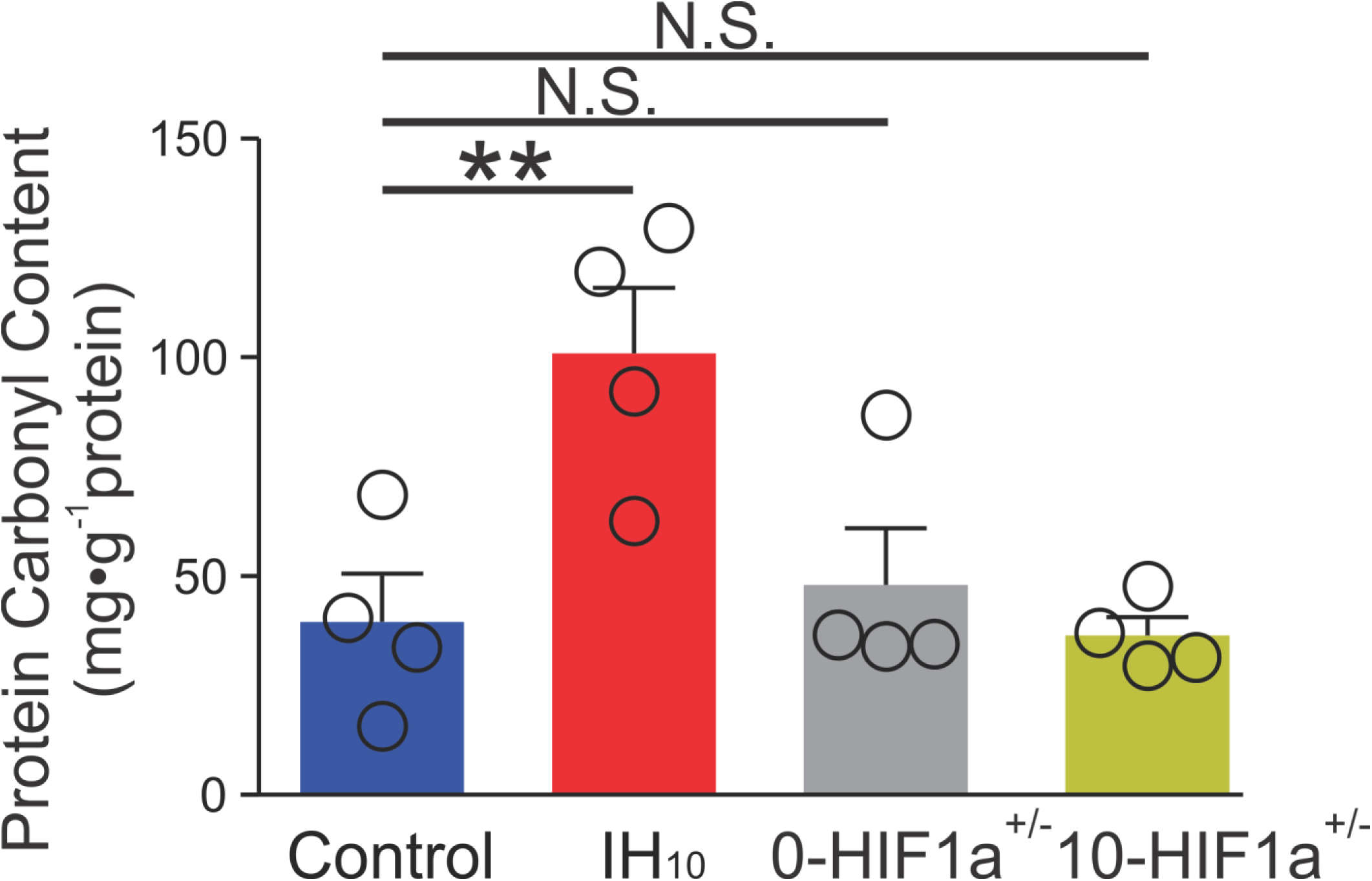
IH increases oxidative stress in the hippocampus. Hippocampal homogenates from control (n=4), IH_10_ (n=4), 0-HIF1a+/- (n=4), and 10-HIF1a+/- (n=4). While IH_10_ displayed elevated protein carbonyl content (P<0.01), protein carbonyl content was not elevated in either 0-HIF1a^+/-^ (P>0.05) or 10-HIF1a^+/-^ (P=0.05). One-way ANOVA comparison was performed followed by a *posthoc* Dunnett’s test. **=P<0.01; N.S.=P>0.05.

**Figure 8.**
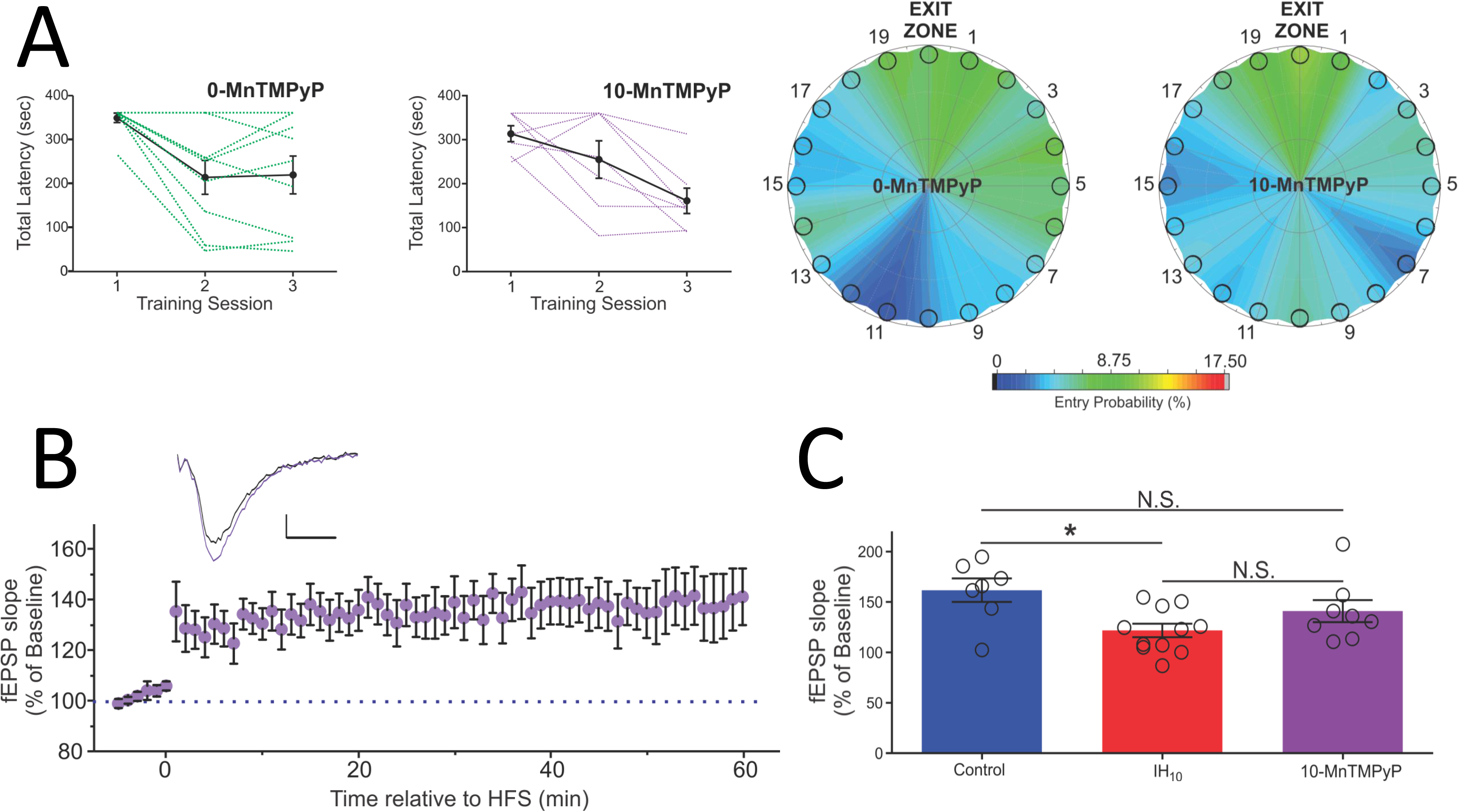
Antioxidant treatment does not fully mitigate the effects of IH on LTP in area CA1. **A**. Animals exposed to MnTMPyP demonstrate similar behaviors following training and probe sessions. Both control (n=9) and IH animals (n=7) had similar latencies during the training period. During the probe trial, both heatmaps demonstrate similar amounts of time spent at the exit zone. **B.** Representative traces of the fEPSP illustrate baseline (black) and 60 mins following HFS (purple trace) in area CA1 from 10-MnTMPyP. Scale bars 0.2 mV/10 ms, **C.** The magnitude of LTP from control, IH_10_, and 10-MnTMPyP was compared. While LTP magnitude in IH_10_ was smaller when compared to control, LTP magnitude from that was not different from either control or IH_10_. Data for control and IH_10_ taken from Figure 6. *=P<0.05, N.S.=P 0.05.

## Discussion

Consistent with previous reports that assess how intermittent hypoxia impacts spatial learning and memory (Wallace and Bucks, 2013; Varga et al., 2014; Gildeh et al., 2016), we observed that IH_10_ disrupted performance in the Barnes maze. This disruption coincided with increased HIF1a, increased oxidative stress, changes in ionotropic glutamate receptors, and impaired LTP. Heterozygosity in HIF1a prevented IH-dependent changes to hippocampal biochemistry, the attenuation of synaptic plasticity in area CA1, and performance deficits in the Barnes maze. Antioxidant administration did not fully attenuate the effects of IH on the performance of wildtype mice in the Barnes maze, or on synaptic plasticity in the hippocampus.

When evaluating behavioral performance in the Barnes maze, both control and IH_10_ groups progressively improved with training, yet prominent differences were present during the probe trial. In addition to the larger primary latency and primary distance traveled following IH, we observed a decrease in the accuracy and precision of locating the exit zone in wildtype mice following IH. These behavioral differences coincided with IH-dependent attenuation of LTP. Our electrophysiological experiments also indicated that IH attenuates the contribution of NMDAr, which would lead to a reduction in NMDAr-dependent Ca^2+^ entry into neurons and diminish the downstream intracellular signaling critical to LTP.

The IH-mediated pro-oxidant condition may have influenced the activity of the NMDAr, as oxidative modulation of the receptor is a well-documented phenomenon (Choi and Lipton, 2000; Lipton et al., 2002; Kumar, 2015; Foster et al., 2017). Alternatively, IH may have downregulated the expression of the functional receptor in the hippocampus. In support of this possibility, IH has also been shown to cause a reduction in NR1, the obligatory subunit for the functional NMDAr, cell density in the cortex and area CA1 (Gozal et al., 2001). Independent of cause, the reduced contribution of the NMDAr appeared to be accompanied by IH-dependent increase in the relative contribution of AMPAr as the fEPSP from control and IH_10_ were similarly stimulated by Mg^2+^-free media yet differentially affected by APV. In cancer cells, the AMPAr subunit proteins GRIA2 and the GRIA3 increases with hypoxia and appears to involve HIF1 signaling (Hu et al., 2014). Thus, our observations indicate that IH-dependent behavioral performance deficits are associated with a reduction in long-term synaptic plasticity and a remodeling of the glutamatergic receptors within the hippocampus.

Employing pharmacological means or IH to enhance HIF1a in the hippocampus correlated with the suppression of LTP in area CA1 (Wall et al., 2014). Similarly, oxidative stress has been implicated as a significant factor that could influence neurophysiological and behavioral outcomes in response to IH (Xu et al., 2004). While HIF1a is commonly considered a pro-survival molecule during hypoxia, HIF1a acts as an upstream factor promoting a pro-oxidant state as it can lead to the expression of pro-oxidant proteins such as NADPH oxidase (Nanduri et al., 2015). Furthermore, when the CCAT/enhancer binding homologous protein (CHOP) is genetically ablated, oxidative stress and HIF1a was not elevated in area CA1 of rodents exposed to IH, suggesting a potential relationship between HIF1a and oxidative stress (Chou et al., 2013). IH did not cause an increase in nuclear HIF1a, or lead to enhanced protein carbonyl content HIF1a^+/-^. These findings are consistent with the idea that IH-dependent HIF1a signaling causes oxidative stress.

No performance deficits caused by IH were observed in HIF1a^+/-^ mice. At the circuit level, IH strengthened glutamatergic transmission as the relationship between current stimulus and the fEPSP, by enhancing contribution of NMDAr and in tissue from 10-HIF1a^+/-^. Combined with our observations in wildtype mice, IH appears to upregulate AMPAr contribution and downregulate NMDAr contribution. While downregulation of NMDAr appears to be dependent on HIF1a, it is unclear whether AMPAr upregulation was HIF1a dependent. Together with our observations in wildtype mice, IH-dependent HIF1a signaling appears to be important for determining the consequences of IH on hippocampal synaptic physiology and hippocampal associated behaviors.

In wildtype mice, IH caused an increase in protein carbonyl content, a measure of oxidative stress within the hippocampus. However, in HIF1a^+/-^ no such increase was observed following IH. Although we did not identify the source of oxidative stress, we demonstrated that antioxidant supplementation using MnTMPyP, to scavenge superoxide anion produced by IH, mitigated many of the IH-mediated changes to behavior and synaptic plasticity. When compared to 0-MnTMPyP, mice receiving MnTMPyP during IH exposure (i.e., 10-MnTMPyP) did not exhibit performance deficits in the Barnes Maze. However, LTP in slices from 10-MnTMPyP was neither similar to LTP recorded in slices from control nor different from LTP evoked in tissue from mice exposed to IH. These findings were unexpected, as antioxidant administration prevented both oxidative stress and behavioral impairments caused by IH (Row et al., 2003). Similarly, MnTMPyP has been shown to mitigate many of the consequences of IH throughout nervous system (Kumar et al., 2006; Peng et al., 2013; Garcia et al., 2016). However, treatment with MnTMPyP may have unique effects in area CA1. Acute incubation in MnTMPyP has been shown to block LTP induction in area CA1 of hippocampal slices (Klann, 1998). This raised the possibility that *in vivo* MnTMPyP treatment could have caused interference with mechanisms common to *in vitro* LTP and spatial learning and memory associated with area CA1. Alternatively, IH may also act independently of reactive oxygen species (ROS) to affect area CA1 and/or other brain networks involved with learning and memory. For example, we have recently demonstrated that IH can cause a ROS-independent expansion of the neuroprogenitor pool involved with adult hippocampal neurogenesis (Khuu et al., 2019). Thus, mechanisms independent of ROS may also be involved in influencing synaptic plasticity, learning, and memory, but this possibility requires further investigation.

In conclusion, our study has demonstrated that IH causes deficits in hippocampal associated behavior, which is correlated with changes in synaptic properties and a weakening in LTP within area CA1 of the hippocampus. We have also shown that HIF1a and oxidative stress appear to be significant factors in dictating these IH-dependent outcomes at the local circuit and behavioral levels. As a result, we have established a working model by which IH-dependent HIF1a signaling acts to cause oxidative stress, remodel glutamatergic synapses, weaken synaptic plasticity, and impair learning and memory.

## Acknowledgements

The authors wish to thank for Dr. N. Prabhakar and Dr. G. Semenza for the provision of the HIF1a^+/-^ mouse line. The authors would also like to thank Dr. N. Prabhakar for the sound advice throughout the course of the study and the preparation of the manuscript. This work was supported by NIH R01 NS10742101 awarded to AJG and a grant from The BSD Office of Diversity & Inclusion at The University of Chicago awarded to AJG. GD was also supported by the University of Chicago Diabetes Research Center (P30 DK020595).

## Author Contribution

AJG conceived and designed experiments; AAC, MAK, JEB, CUN, GD and AJG performed experiments, and conducted analyses; AAC, MAK, and AJG wrote and/or revised the manuscript; AJG provided unpublished reagents and analytic tools.

## Conflict of Interest

The authors declare no competing financial interests.

## Notes

#### Summary of Updates

Minor Revisions to text and references were made for clarity. Figure 5 revised

